# Burst firing optimizes invariant coding of natural communication signals by electrosensory neural populations

**DOI:** 10.1101/2024.09.09.611671

**Authors:** Michael G. Metzen, Amin Akhshi, Pouya Bashivan, Anmar Khadra, Maurice J. Chacron

**Affiliations:** Department of Physiology, McGill University, Montreal, QC, Canada

**Keywords:** burst firing, population coding, weakly electric fish, neural circuits

## Abstract

Accurate perception of objects within the environment independent of context is essential for an organism’s survival. While neurons that respond in an invariant manner to identity-preserving transformations of objects are thought to provide a neural correlate of context-independent perception, how these emerge in the brain remains poorly understood. Here we demonstrate that burst firing in neural populations can give rise to an invariant representation of highly heterogeneous natural communication stimuli. Multi-unit recordings from central sensory neural populations showed that considering burst spike trains led to invariant representations at the population but not the single neuron level. Computational modeling further revealed that optimal invariance is achieved for levels of burst firing seen experimentally. Taken together, our results demonstrate a novel function for burst firing towards establishing invariant representations of sensory input in neural populations.

## Introduction

Every organism must accurately recognize objects within its sensory environment independent of context to survive. Towards this end, neural representations that are invariant to identity preserving transformations of sensory input have been found across sensory systems, including visual [1-7]; auditory [8, 9] and olfactory [10] systems. Such representations are thought to progressively emerge across successive brain areas by making neurons more and more invariant to identity preserving transformations by means of so-called “OR” operations (i.e., respond when stimulus A or stimulus B is presented). However, the mechanisms that mediate such progressive emergence remain poorly understood.

The electrosensory system of gymnotiform wave-type weakly electric fish benefits from well-characterized anatomy and simple natural stimuli that can easily be mimicked in the laboratory and shares many similarities with the mammalian visual, auditory, and vestibular systems [11, 12]. These fish continuously produce an electric field around their body through the electric organ discharge (EOD) and sense amplitude modulations of this field through an array of peripheral electroreceptor afferents that synapse onto pyramidal cells within the electrosensory lateral line lobe (ELL). ELL pyramidal cells in turn project to the midbrain torus semicircularis (TS) and indirectly to higher brain areas mediating behavior [13]. Natural electrosensory stimuli comprise those encountered during social interactions. Specifically, conspecifics can communicate with one another through transient increases in EOD frequency called “chirps” [14-20]. Chirps with different attributes (i.e., duration, amplitude) give rise to widely different stimulus waveforms.

Previous work has characterized the responses of electroreceptor afferents [21-25], ELL pyramidal cells [15, 26-31], and downstream neurons [32, 33] to chirps. Notably, while the firing activities of single electroreceptor afferents faithfully follow the chirp stimulus waveforms, the activities of electroreceptor afferent populations are transiently synchronized during the chirp, providing an invariant representation [25]. Recordings from single ELL and TS neurons reveal progressively more invariant responses, thereby explaining the animal’s behavioral responses [25, 31, 34].

ELL pyramidal cells display an active burst mechanism relying on a somato-dendritic interaction via spike backpropagation [35-38]. Specifically, bursts are initiated when action potentials backpropagate from the soma to the dendritic tree, which triggers dendritic spikes that propagate back to the soma, thereby giving rise to a depolarizing afterpotential. This “ping-pong” mechanism continues until the interspike interval becomes smaller than the dendritic refractory period, which leads to dendritic failure and burst termination [35, 37, 39-42]. While previous studies have demonstrated multiple functions for such burst firing including coding of low frequency stimuli [43, 44] and of looming objects [45], how such burst firing affects response invariance to natural electrocommunication stimuli remains explored to date.

## Results

The goal of this study was to investigate how burst firing impacts population coding of natural electrocommunication stimuli by ELL pyramidal cells. To do so, simultaneous recordings from ELL neural populations were collected using Neuropixels probes (Fig. 1a). Stimuli consisted of natural electrocommunication signals during social interactions (Fig. 1b). These consisted of waveforms resulting from one individual fish (i.e., the “emitter fish) that transiently increases its EOD frequency (i.e., “amplitude”) for a short duration, causing a phase reset of the beat (Fig. 1b). Varying parameters such as amplitude, duration, and the phase of the beat at the onset of the chirp results in very different (i.e., heterogeneous) stimulus waveforms (Fig. 1c). We used the correlation coefficient to quantify similarity between these different waveforms (see Methods). Overall, heterogeneity was reflected in the wide range of correlation coefficients obtained when visualized using the correlation matrix (Fig. 1d). Single ELL pyramidal cells displayed burst firing in response to natural electrocommunication stimuli, as evidenced by their bimodal ISI distribution (supplementary Fig. 1a) and consistent with previous results [29]. We used a standard methodology to separate spikes into two groups: those that belong to bursts (i.e., the burst spike train) and those that did not (i.e., the isolated spike train; supplementary Fig. 1a) using the trough between the two modes as a burst threshold (see Methods).

**Figure 1:**
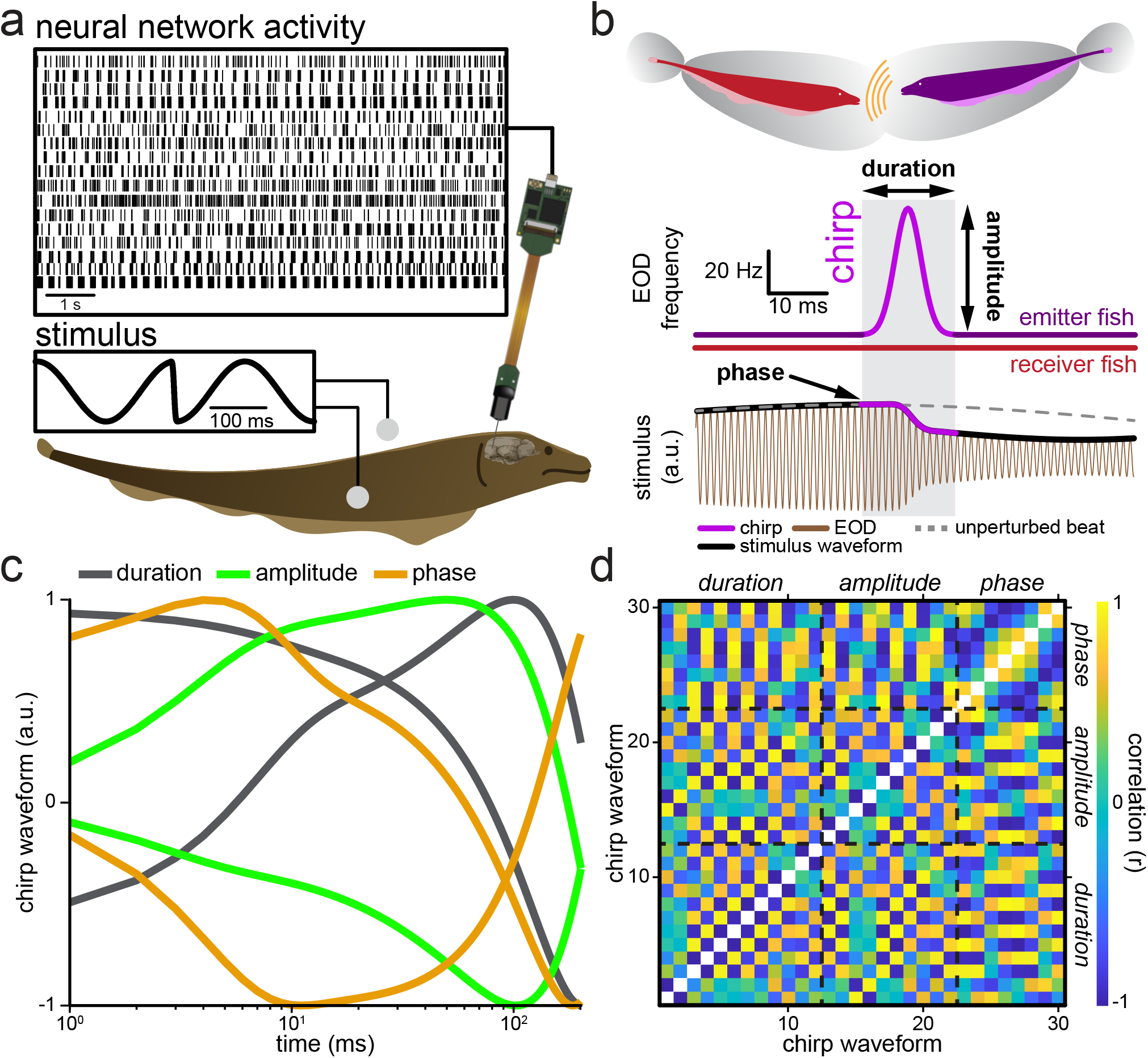
Experimental setup and natural electrocommunication stimuli. (a) Schematic showing the experimental setup. A fish is placed in an otherwise empty tank and is stimulated (middle left) while neuronal activity (top) is recorded using a Neuropixels probe. **(b)** During a natural electrocommunication stimulus event (i.e., a “chirp”), the emitter fish’s (top, purple) EOD frequency (middle, blue trace) is transiently increased to a maximum value for a brief duration (light purple, shaded box) while the receiver fish’s (top, red) EOD frequency (middle, red trace) remains constant. This can be characterized by the duration and amplitude of the frequency excursion. The chirp results in a phase advance of the beat (i.e., the beat will reach its maximum value earlier than if no chirp occurred; compare solid black with dashed gray). The chirp is shown in light purple. **(c)** Examples of natural electrocommunication stimulus waveforms obtained by varying different parameters (gray: chirp duration; light green: chirp amplitude; orange: chirp phase). **(d)** Correlation coefficient matrix for all 30 chirp stimulus waveforms considered in this study.

### Burst firing mediates invariant population coding of natural electrocommunication stimuli

To investigate how burst firing impacts population coding of natural electrocommunication stimuli, we separated the full spike trains of the recorded ELL pyramidal cells into burst and isolated spike trains based on the burst threshold. We note that our methodology considered the structure of bursts (e.g., the number of spikes within each burst), as opposed to treating them as “events”.

The time-varying network activity was obtained by summing across neurons either of the full spike trains, burst spike trains, or isolated spike trains. Overall, the network activities in response to different chirp stimulus waveforms obtained when considering the burst spike trains all consisted of an initial rise following stimulus onset (Fig. 2a, compare red traces within the gray bands) and thus overlapped with one another when superimposed (Fig. 2b, compare red traces). In contrast, network activities obtained when considering either the full or isolated spike trains were more heterogeneous (Fig. 2a, compare black and blue traces) and thus did not overlap when superimposed (Fig. 2b, upper panel, compare black and blue traces). By quantifying the similarity between network activities in response to different stimulus waveforms using the correlation coefficient (see Methods), we found that the correlation coefficients obtained from burst spike trains were more positive and significantly higher than those obtained when considering either the full or isolated spike trains (Fig. 2b, bottom panels; Fig. 2c; one-way ANOVA, df = 2; F = 89.88; p_full,burst_ = 1.21*10^−19^; p_burst,iso_ = 6.08*10^−36^; Bonferroni corrected). Based on this, we conclude that network activities obtained when considering burst spike trains provide a more invariant representation of natural electrocommunication stimuli. This result was robust, as higher correlation values were obtained for a wide range of burst thresholds (supplementary Fig. 1b). Interestingly, similar results were obtained when randomly shuffling responses with respect to stimulus trials, which removes noise correlations (supplementary Fig. 2).

**Figure 2:**
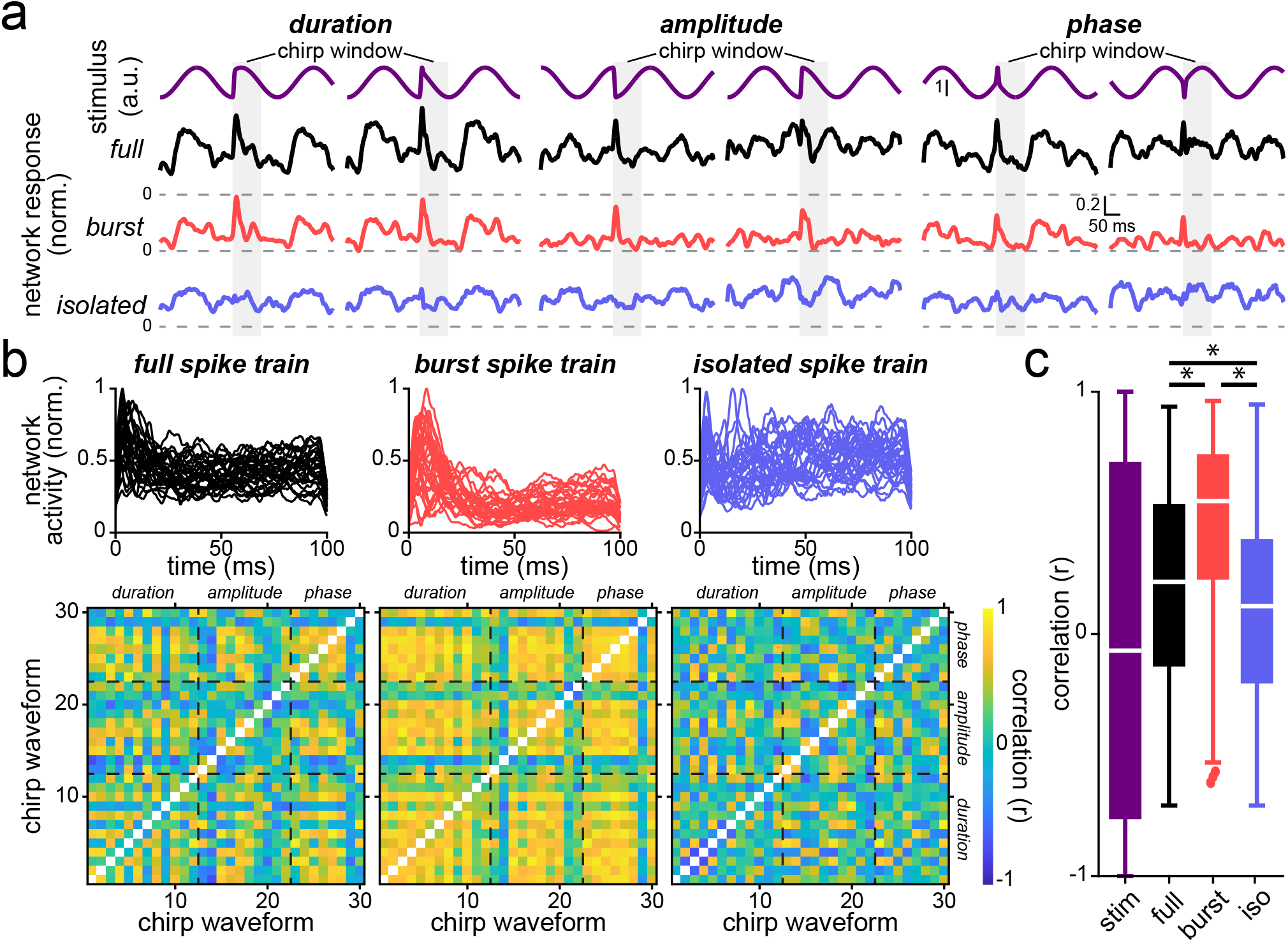
Burst firing increases neural population responses to natural electrocommunication stimuli. **(a)** Examples network activities in response to different chirp waveforms obtained when varying chirp duration (top left, purple), chirp amplitude (top middle, purple) and chirp phase (top right, purple). Shown are the activities obtained using the full (i.e., all spikes, black; middle top), burst (red, middle bottom), and isolated (blue, bottom) spike trains, respectively. Note that the responses of the burst spike trains were more similar to one another (i.e., are more invariant) during the chirp window (gray bands) for different chirp waveforms than either all or isolated spike trains. **(b)** *Top*: Superimposed network activity traces in response to all chirp stimuli when considering either the full (black, left), burst (red, middle) and isolated spike trains (blue, right). *Bottom*: Correlation matrices computed during the chirp time window of network activities when considering the full (left), burst (middle) and isolated spike trains (right). **(c)** Whisker-box plots of correlation coefficients obtained for the stimulus waveforms (purple), as well as for network activities when considering either all (black), burst (red) and isolated (blue) spikes. Correlation coefficients were significantly higher when considering burst spike trains than when considering the full or isolated spike trains (one-way ANOVA, df = 2; F = 89.88; p_full,burst_ = 1.21*10^−19^; p_full,iso_ = 7.85*10^−04^; p_burst,iso_ = 6.08*10^−36^; Bonferroni corrected).

### Invariant coding of natural electrocommunication stimuli by burst firing is seen at the population but not the single neuron level

Perhaps the simplest explanation for our results is that burst firing provides a more invariant representation of natural electrocommunication stimuli at the single neuron level. To test this hypothesis, we instead considered the firing activities of single neurons in response to different chirp stimulus waveforms. Overall, we found that these did not overlap to a great extent when superimposed (Fig. 3a, left upper panel) as quantified by low correlation coefficient values (Fig. 3a, lower left panel; one-way ANOVA, df = 2; F = 48.41; p_full,burst_ = 4.93*10^−21^; p_full,iso_ = 0.002; p_burst,iso_ = 1.24*10^−9^; Bonferroni corrected). Further analysis revealed that this is because the responses of single ELL neurons to different chirp stimulus waveforms were highly heterogeneous when considering either the full, burst or isolated spike trains (supplementary Fig. 3). Taken together, these results indicate that the invariant representation of natural electrocommunication stimuli, derived from network activity of burst spike trains, is not a single neuron property but rather a consequence of population level coding.

**Figure 3:**
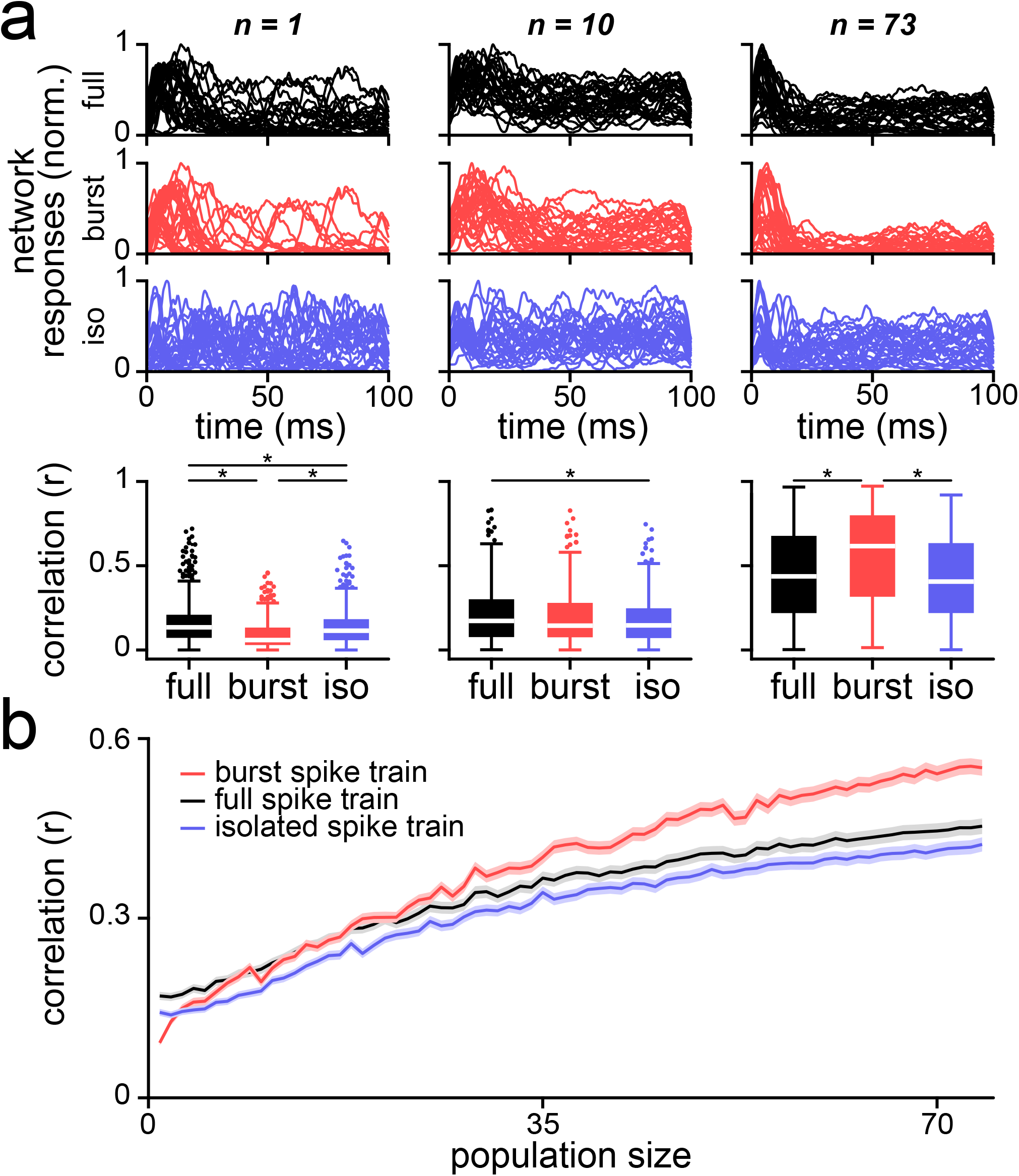
Enhanced response invariance due to burst firing is seen for populations but not single neurons. **(a)** *Top*: S single neuron (left), populations of n = 10 neurons (middle), and populations of n = 73 neurons (right) activities in response to all chirp stimuli when considering all (black), burst (red) and isolated spike trains (blue). Note that each line shows the activity in response to a given stimulus waveform. *Bottom*: Whisker-box plots showing correlation coefficients obtained for either single neurons (left), populations of n = 10 neurons (middle), and populations of n = 73 neurons (right) when considering all (black), burst (red) and isolated (blue) spikes. For single neurons, correlation coefficients for bursts were significantly lower than those obtained for either all or isolated spike trains (one-way ANOVA, df = 2; F = 48.41; p_full,burst_ = 4.93*10^−21^; p_full,iso_ = 0.002; p_burst,iso_ = 1.24*10^−9^; Bonferroni corrected). For populations of n = 10 neurons, correlation coefficients obtained for the full, burst and isolated spike trains were in general not significantly different from one another (one-way ANOVA, df = 2; F = 5.69; p_full,burst_ = 0.16; p_full,iso_ = 0.002; p_burst,iso_ = 0.47; Bonferroni corrected). **(b)**. Correlation coefficients as a function of population size when considering the full (black), burst (red) and isolated spike trains (blue). The error bands show 1 SEM.

To further test this hypothesis, we next systematically assessed the network activity of ELL pyramidal cells in response to different chirp stimulus waveforms at two different population sizes of n = 10 (Fig. 3a, middle panels) and n = 73 (Fig. 3a, right panels). Our results revealed that, as the number of cells in the population increased, the network activities obtained when considering burst spike trains progressively overlapped more and more with one another (Fig. 3a, compare red traces in middle and right panels) as quantified by higher correlation values (Fig. 3a, lower middle and right panels). Greater overlap between the network activities in response to different chirp stimulus waveforms was also observed in the two cases of the full and isolated spike trains when the population size increased (Fig. 3a, compare black and blue traces in middle and right panels). In fact, for a population size of n = 10, correlation values obtained when considering either the full, burst, or isolated spike trains were not significantly different from one another, except for the full and isolated spike trains (Fig. 3a, bottom middle panel; one-way ANOVA, df = 2; F = 5.69; p_full,burst_ = 0.16; p_full,iso_ = 0.002; p_burst,iso_ = 0.47; Bonferroni corrected). In contrast, for a population size of n = 73, network activities obtained when considering burst spike trains better overlapped with one another than those obtained from either the full or isolated spike trains (Fig. 3a, upper right panels, compare red, black, and blue traces) as quantified by significantly higher correlation coefficient values (Fig. 3a, bottom right panel; one-way ANOVA, df = 2; F = 31.17; p_full,burst_ = 3.44*10^−8^; p_full,iso_ = 0.21; p_burst,iso_ = 2.23*10^−13^; Bonferroni corrected). Indeed, correlation coefficients obtained when considering burst spike trains increased at a higher rate as a function of population size than those obtained when considering either the full or isolated spike trains (Fig. 3b; compare red, black, and blue traces). This increase in correlation coefficient values obtained when considering burst spike trains initially started from lower values but diverged above those obtained using either the full or isolated spike trains as population size increased (Fig. 3b; compare red, black, and blue traces).

### A two-compartment model of ELL pyramidal cell activity displays invariant coding of natural electrocommunication stimuli

So far, our results have shown that the network activities in response to different chirp stimulus waveforms obtained when considering burst spike trains were more similar (i.e., invariant) than those obtained when considering either the full or isolated spike trains. Further analysis revealed that this effect is not seen at the single neuron level but emerges when the population size is sufficiently large (i.e. > ∼30). In order to gain further understanding as to why burst firing provides a more invariant representation at the population level, we used a computational model of ELL pyramidal cell activity (see Methods). The model consisted of somatic and dendritic compartments each with ionic currents necessary to reproduce burst firing displayed by ELL pyramidal cells (Fig. 4a; see Methods). The model was fit to experimentally recorded spike trains from each ELL pyramidal cell in our dataset using maximum likelihood and the activities from the model neurons in response to different chirp stimulus waveforms were then analyzed in the same manner as our experimental data (see Methods).

**Figure 4:**
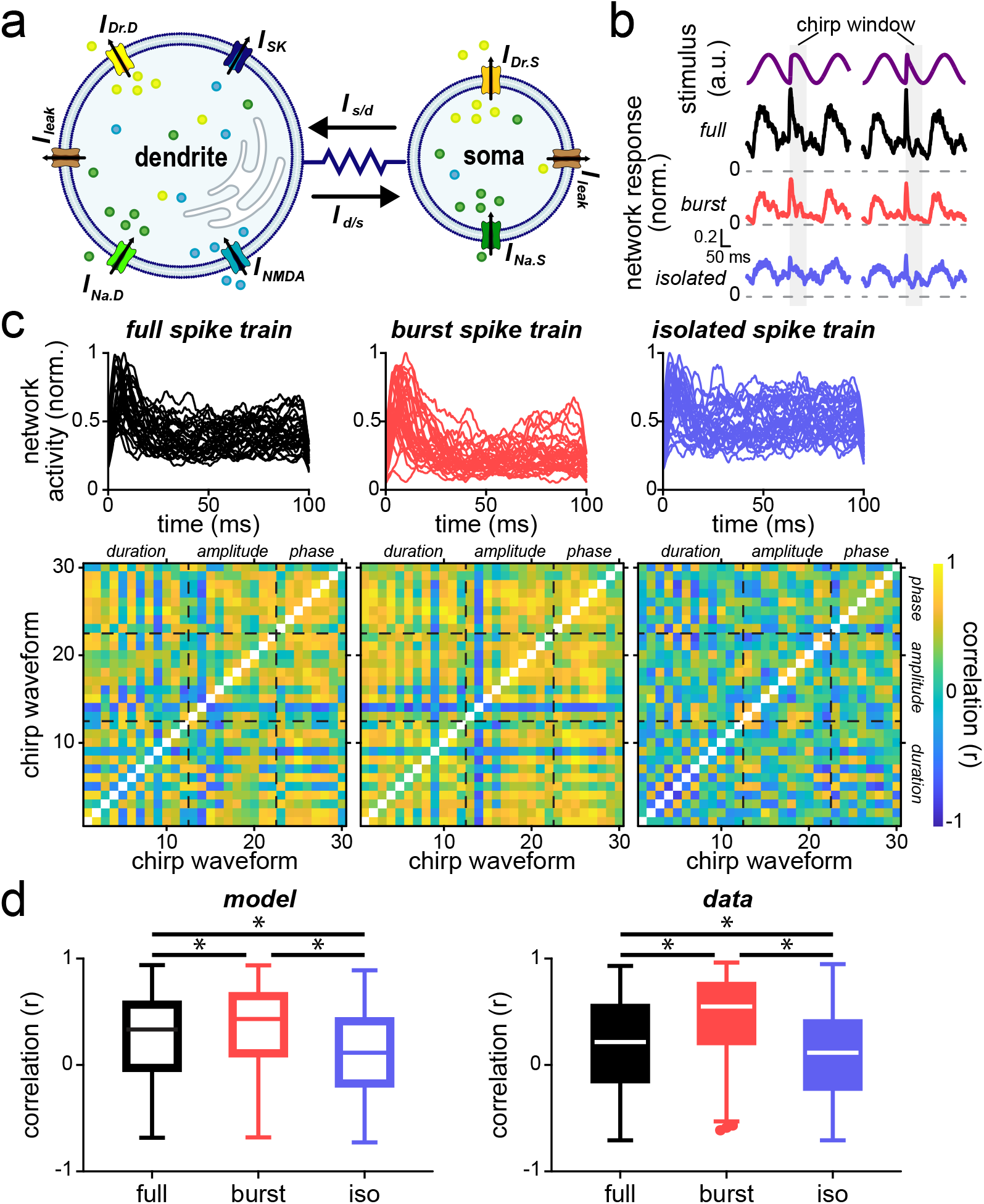
A two-compartment model of ELL pyramidal cell activity shows enhanced response invariance to natural electrocommunication stimuli when considering burst spike trains. **(a)** Schematic of the model showing the somatic and dendritic compartments with sodium, delayed rectifier potassium, leak, and NMDA and SK currents. **(b)** Model network activity in response to several chirp waveforms (top, purple) using the full (i.e., all spikes, black; middle top), burst (red, middle bottom) and isolated (blue, bottom) spike trains, respectively. **(c)** *Top*: Superimposed network activity traces from the model in response to all chirp stimuli when considering either the full (black, left), burst (red, middle) and isolated spike trains (blue, right). *Bottom*: Correlation matrices of network activities when considering the full (left), burst (middle) and isolated spike trains (right). **(d)** Whisker-box plots of correlation coefficients obtained for network activities for the model (left, hollow) and for experimental data (right, solid) when considering either the full (black), burst (red), and isolated (blue) spike trains. Correlation coefficients were significantly higher when considering burst spike trains than when considering the full or isolated spike trains generated by the model (one-way ANOVA, df = 2; F = 89.88; p_full,burst_ = 1.21*10^−19^; p_full,iso_ = 7.85*10^−04^; p_burst,iso_ = 6.08*10^−36^; Bonferroni corrected). There was an excellent match between model and data (one-way ANOVA, df = 5; F = 17.75; p_full_ = 0.15; p_burst_ = 0.23; p_iso_ = 0.99; Tuckey-Kramer corrected).

Overall, we found that the model could consistently reproduce our experimental results. Indeed, the network activities in response to different chirp waveforms obtained when considering the burst spike trains all consisted of an initial rise following stimulus onset (Fig. 4b, compare red traces within the gray bands) and thus overlapped with one another when superimposed (Fig. 4c, compare red traces). In contrast, network activities obtained when considering either the full or isolated spike trains were more heterogeneous (Fig.4b, upper panels, compare black and blue traces) and thus did not overlap to the same extent as those for burst spike trains when superimposed (Fig. 4c, upper panels, compare red and blue traces). Correlation coefficient values were on average more positive and thus significantly higher when considering burst spike trains than when considering either the full or isolated spike trains (Fig. 4c, bottom panels; Fig. 4d; one-way ANOVA, df = 2; F = 46.46; p_full,burst_ = 0.001; p_full,iso_ = 7.80*10^−9^; p_burst,iso_ = 2.13*10^−20^; Bonferroni corrected). Importantly, correlation coefficient values from the model obtained were not significantly different than those obtained experimentally when considering either the full, burst, or isolated spike trains (one-way ANOVA, df = 5; F = 17.75; p_full_ = 0.15; p_burst_ = 0.23; p_iso_ = 0.99; Tuckey-Kramer corrected).

### Invariant population coding of natural electrocommunication stimuli by burst firing is optimal for intermediate levels of burst firing

We next varied model parameters and investigated their effect on invariant population coding by burst firing. In particular, we varied the level of burst firing in the model by changing the value of the maximum dendritic sodium conductance *g*_*NaD*_ (Fig. 5a). We chose this parameter because, as mentioned above, burst firing in ELL pyramidal cells is dependent on dendritic spiking. Thus, varying *g*_*NaD*_ will give rise to different levels of bursting in our model as quantified by the burst fraction (i.e., the fraction of ISIs below threshold; see Methods). We found that invariant population coding of natural electrocommunication stimuli by burst firing was compromised when the model displayed much lower (higher) levels of burst firing corresponding to smaller (larger) values of *g*_*NaD*_ (supplementary Figs. 4, 5, respectively). We next systematically varied *g*_*NaD*_ and plotted the correlation coefficient for bursts as a function of the burst fraction, which was varied by changing the burst threshold (Fig. 5b; see Methods). For low *g*_*NaD*_ values (i.e., low levels of burst firing), maximum correlation was obtained for highest burst fractions. In contrast, for higher *g*_*NaD*_ values, the correlation was maximum for intermediate values of burst fraction.

**Figure 5:**
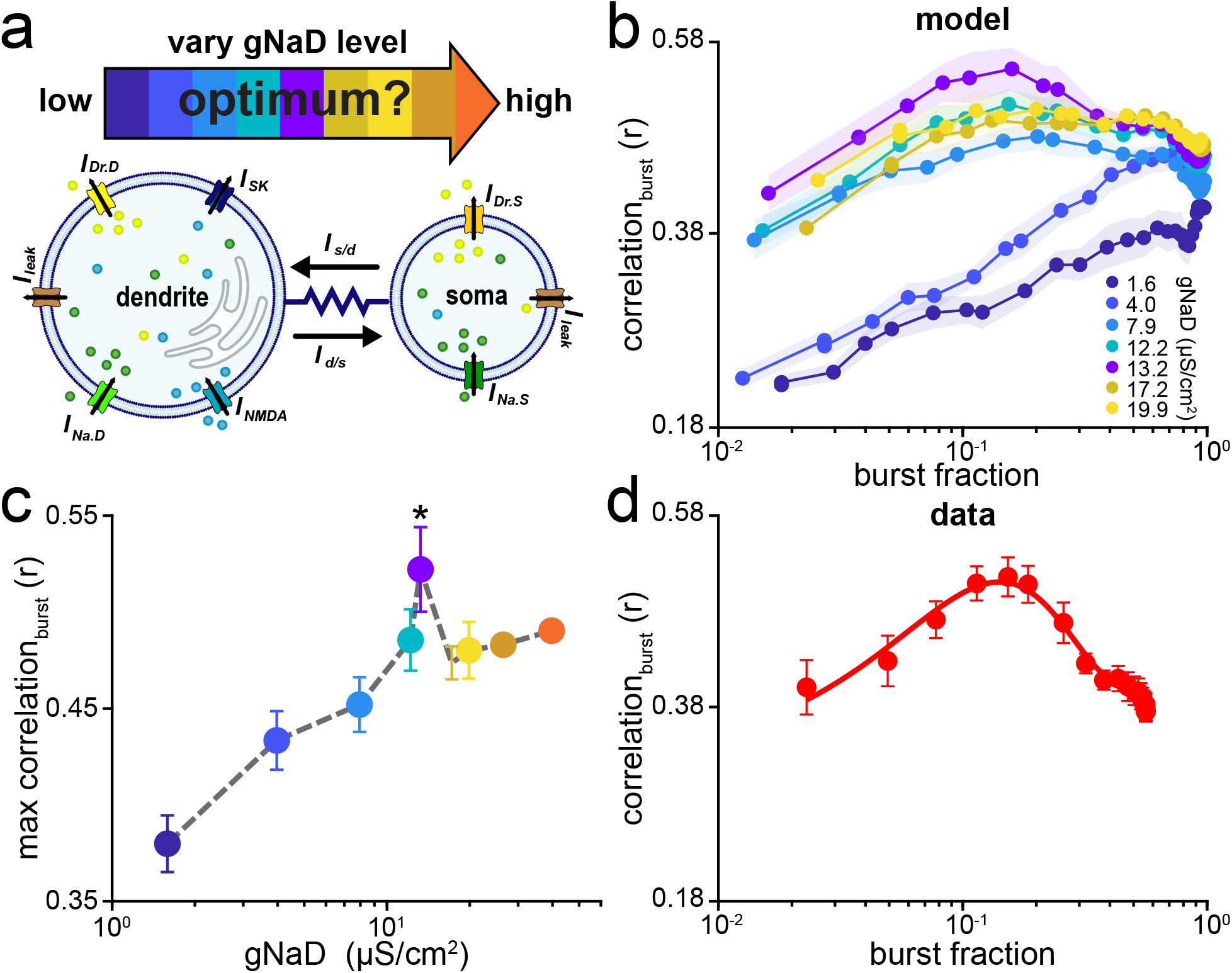
Invariant representations of natural electrocommunication stimuli is maximized for intermediate levels of burst firing. **(a)** Model schematic showing that the level of burst firing can be tuned by varying the maximum dendritic sodium conductance *g*_*NaD*_. **(b)** Correlation as a function of burst fraction obtained for the model for different values of *g*_*NaD*_. **(c)** Maximum correlation for burst spike trains as a function of *g*_*NaD*_ level. The optimal correlation was significantly higher than the other values (one-way ANOVA, df = 8; F = 233.60; p < 3.06*10^−17^; Bonferroni corrected) **(d)** Correlation as a function of burst fraction for experimental data. Note the similarity with the curve obtained from the model for the optimal value of *g*_*NaD*_.

Importantly, the highest correlation was obtained for values of *g*_*NaD*_ that gave rise to levels of burst firing similar to those seen experimentally (Fig. 5c). Our model thus makes two important predictions: 1) there is an optimal level of *g*_*NaD*_, and 2) maximum response correlation for burst firing is obtained for a certain level of population burst fraction. We tested prediction 2) by plotting the correlation values from experimental data as a function of burst fraction. We found that correlation was indeed maximal for an intermediate value of burst fraction, which corresponded to the value obtained when separating burst from isolated spikes using the trough of the bimodal ISI distribution (Fig. 5d). These results suggest that burst firing is regulated in ELL pyramidal cells to optimize invariant population coding of natural electrocommunication stimuli.

### ELL pyramidal cells display no trade-off between discriminability and invariance

Finally, we investigated whether invariant population coding by burst firing occurred at the expense of stimulus discriminability. To do so, we quantified the performance of a classifier at assigning a given network activity as being caused by a given chirp stimulus waveform (see Methods). We note that this analysis has been previously used across sensory systems [32, 46-48]. Specifically, template network activities (i.e., the network activity obtained for one trial) were assigned to each chirp stimulus waveform and the remaining activities (i.e., the network activities obtained for the remaining 39 trials) were then compared to each template using a distance metric. The observed activity was assigned to the chirp stimulus waveform whose template was closest as determined by the minimum distance. The discrimination performance was computed from the confusion matrix whose element *ij* gives the conditional probability that a response generated by stimulus *i* is classified as being generated by stimulus *j*. The diagonal elements represent correct assignments while off-diagonal elements instead represent incorrect assignments. Thus, if our classifier displayed 100% correct performance, then the diagonal of the confusion matrix would be unity with all off-diagonal elements zero.

Overall, discrimination performance was highest for network activities obtained when considering the full spike train (Fig. 6a; Fig. 6b, black; one-way ANOVA; df = 2; F = 5.44*10^+3^; p_full,burst_ = 4.22*10^−158^; p_full,iso_ = 7.24*10^−65^; p_burst,iso_ = 2.45*10^−184^; Bonferroni corrected). Further analysis revealed that this was because of isolated and not burst spike trains. Indeed, higher discrimination performances were observed when considering isolated spike trains as opposed to burst spike trains (Fig. 6a; Fig. 6b, left panel, compare blue and red). This contrasts to network activities obtained when considering burst spike trains showing greater invariance than those obtained when considering isolated spike trains as quantified by correlation (Fig. 6b, right panel, red; one-way ANOVA; df = 2; F = 9.98*10^3^; p_full,burst_ = 1.49*10^−215^; p_full,iso_ = 1.10*10^−172^; p_burst,iso_ = 9.87*10^−124^; Bonferroni corrected). Thus, our results show that ELL pyramidal cell populations transmit information that can be used to both reliably identify natural electrocommunication stimuli while maximizing invariance.

**Figure 6:**
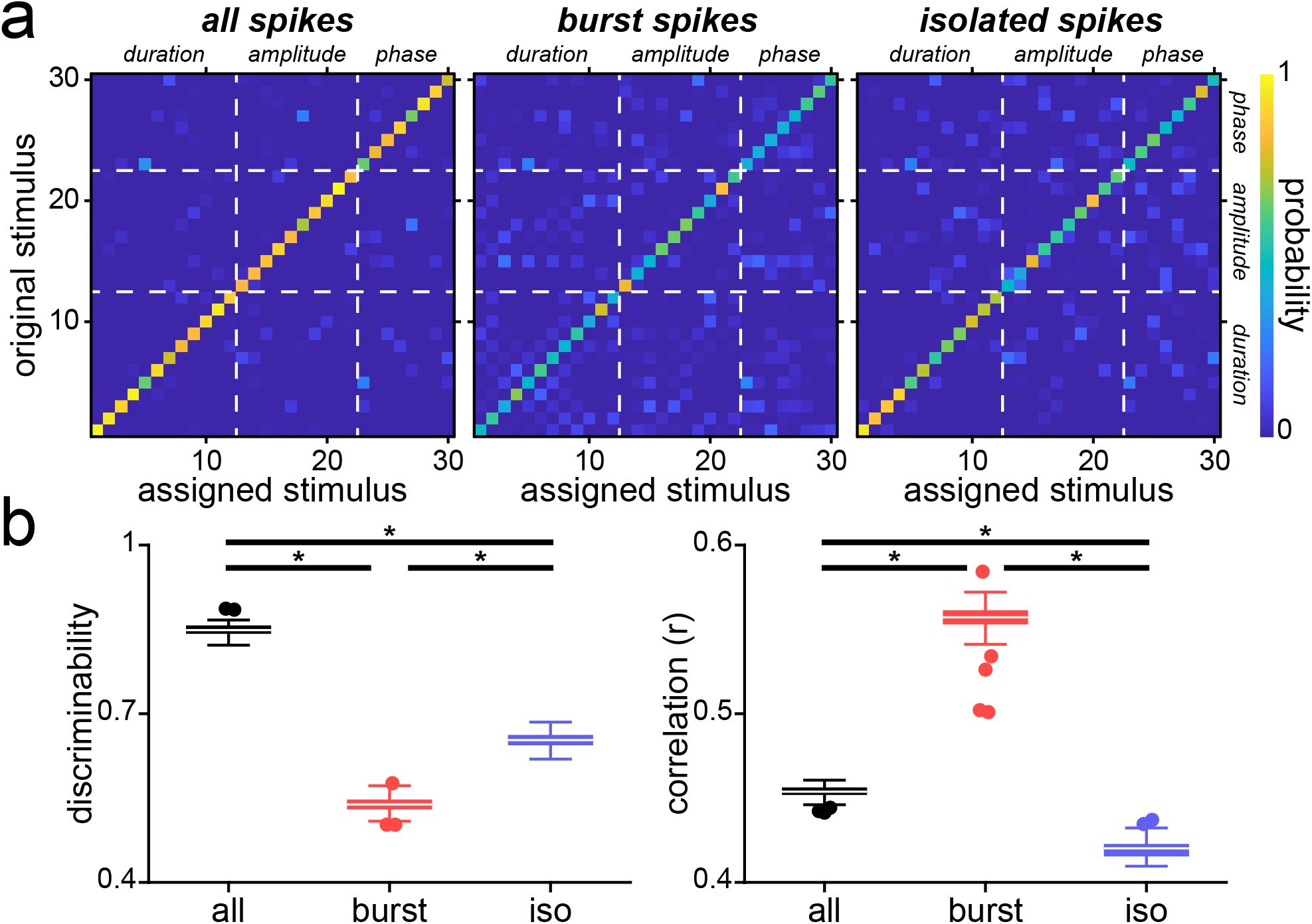
Discriminability and invariance are not mutually exclusive. **(a)** Confusion matrices showing the probability of classifying the population response to be caused by stimulus *i* when it was actually caused by stimulus *j* computed from metric-space analysis using van Rossum measure with a timescale of *τ* = 3 ms for the full (left), burst (middle) and isolated spike trains (right). The white dashed lines indicate borders between blocks of stimuli for which a given attribute (e.g., duration, amplitude, phase) was varied. **(b)** Discrimination performance was significantly higher when considering the full spike train (left; one-way ANOVA; df = 2; F = 5.44*10^+3^; p_full,burst_ = 4.22*10^−158^; p_full,iso_ = 7.24*10^−65^; p_burst,iso_ = 2.45*10^−184^; Bonferroni corrected). This was different for the correlation, where burst spike trains outperformed the full and isolated spike trains (right; one-way ANOVA; df = 2; F = 9.98*10^+3^; p_full,burst_ = 1.49*10^−215^; p_full,iso_ = 1.10*10^−172^; p_burst,iso_ = 9.87*10^−124^; Bonferroni corrected).

## Discussion

### Summary of results

We investigated how burst firing impacted population coding of natural electrocommunication stimuli by ELL pyramidal cells. We found that network activities in response to different stimulus waveforms were more similar to one another (i.e., were invariant) when considering burst spike trains than when considering either the full or isolated spike trains. Further analysis revealed that such population invariance was seen at the population rather than at the single neuron level. To gain more understanding as to how burst firing can give rise to population responses that are invariant to natural electrocommunication stimuli, we used a computational model of burst firing. We found that our model accurately reproduced experimental results. Further, model simulations revealed that invariance was optimal for intermediate levels of burst firing similar to those seen experimentally. Interestingly, network activities obtained when considering isolated spike trains allowed for better discrimination of chirp stimulus waveforms than bursts.

### Invariant population coding: a novel function for burst firing

Our results show that burst firing provides, at the population level, a more invariant representation of natural electrocommunication stimuli than either the full or isolated spike trains. This is, to our knowledge, a novel function for burst firing. Indeed, previous studies have shown a variety of functions for burst firing including enhanced detection of specific stimulus features [45, 49-53], increasing reliability of synaptic transmission [54], inducing synaptic plasticity [55], encoding of low frequency stimulus features [43, 44, 56, 57], prediction errors [58, 59], and cortical spindle generation [60]. The function uncovered in the current study, however, greatly differs from enhanced feature detection. Indeed, our results show that similar patterns of burst firing are elicited at the population level by different stimulus waveforms, in contrast to previous results that have shown bursts in single neurons are more reliably elicited by specific stimulus features (e.g., upstrokes) [49-54].

### Regulation of burst firing by ELL pyramidal cell populations

Our results suggest that the level of burst firing displayed by ELL pyramidal cells maximizes invariant population coding of natural electrocommunication stimuli. Previous studies have shown that burst firing in ELL pyramidal cells is regulated by multiple mechanisms including small conductance calcium activated potassium channels [61, 62] as well as neuromodulators such as acetylcholine [63] and serotonin [64-66] (see [67-69] for review). Burst firing in ELL pyramidal cells is furthermore regulated by descending pathways [70].

### Decoding invariant representations of natural electrocommunication stimuli

Information transmitted by neural populations within a given brain area is only useful to an organism if it is decoded downstream. An important question is then: are burst spike trains from ELL neural populations decoded by their downstream TS neurons? Previous studies have shown that TS neurons display subthreshold membrane conductances that allow for increased detection of coincident synaptic input [71, 72]. Interestingly, previous studies have shown that some single TS neurons respond in an invariant fashion to natural electrocommunication stimuli [25, 32-34, 73, 74]. We hypothesize that these TS neurons respond selectively to burst firing from afferent ELL neural populations and that, as such, burst firing is essential towards building an invariant representation of natural electrocommunication stimuli within the electrosensory brain. Further studies are needed to test this hypothesis.

### Parallel electrosensory pathways allow for simultaneous detection and identification of natural electrocommunication stimuli

Indeed, while bursts provided an invariant representation of natural electrocommunication stimuli, isolated spikes instead could be used to discriminate between different waveforms. Thus, it is possible for the electrosensory brain to get information allowing for simultaneous detection and identification of chirp waveforms through decoding information transmitted by ELL neural populations by considering either bursts or the full spike trains. As mentioned above, the fact that some TS neurons respond in an invariant fashion to natural electrocommunication stimuli [25, 31, 33] suggest that this is indeed the case. The activities of these neurons provide reliable detection of the chirp waveform independently of its attributes and most likely mediate the animal’s behavioral responses (i.e., “echo responses”) [75]. However, other TS neurons do not respond to natural electrocommunication stimuli in an invariant fashion. Rather, their responses resemble that of ELL neurons (i.e., are “ELL-like”) and, as such, their activity can be used to reliably discriminate between different stimulus waveforms [32, 73]. This suggests that information regarding chirp identity is transmitted by TS neural subpopulations to higher brain areas. Interestingly, our results show that considering the full spike trains led to network activities that could better discriminate between different chirp stimulus waveforms than when considering the isolated spike trains. This suggests that bursts and isolated spikes act in a synergistic manner to provide information regarding chirp identity. We hypothesize that this is because we considered the full structure of the burst spike train, which can provide some information as to chirp identity (e.g., through changes in the number of spikes per burst). Further studies are needed to test whether information as to chirp identity is decoded downstream of TS to mediate perception.

Finally, we note that our results in no way imply that burst firing cannot be used to discriminate between stimuli associated with different behavioral contexts (e.g., those caused by prey as opposed to those caused by conspecifics). Indeed, previous studies have shown that burst firing could be used to gain information as to object location and direction of movement [43, 45, 76]. One interesting possibility is that bursts could be used to discriminate between stimuli associated with different behavioral contexts, while isolated spikes are instead used to discriminate between stimuli within a given context. Such parallel coding of “global” as opposed to “local” features has been demonstrated elsewhere [77].

### Applicability to other systems

It is likely that our results extend to other systems. First, burst firing has been observed ubiquitously across brain areas (see [78, 79] for review). Moreover, invariance develops progressively across successive brain areas and systems, including visual [1-7]; auditory: [8, 9] and olfactory [10]) systems. Importantly however, these studies only have considered the full spike trains as opposed to specific spike patterns such as bursts. We showed in this study that burst firing played an important role towards developing invariant representations of sensory input in these systems as well.

## Methods

### Ethics statement

All animal procedures were approved by McGill University’s animal care committee (# 5285) and were performed in accordance with the guidelines of the Canadian Council on Animal Care. All fish were immobilized by an intramuscular injection of 0.1 – 0.5 mg of tubocurarine prior to experimentation.

### Animals

The wave-type weakly electric fish *Apteronotus leptorhynchus* (N = 2) of either sex was used exclusively in this study. Animals of either sex were purchased from tropical fish suppliers and were housed in groups (2 – 10) at controlled water temperatures (26 – 29°C) and conductivities (300 – 800 µS*cm^-1^) according to published guidelines [80].

### Surgery

Surgical procedures have been described in detail previously [61, 81, 82]. Briefly, 0.1-0.5 mg of tubocurarine (Sigma) was injected intramuscularly to immobilize the animals for electrophysiology experiments. The animals were then transferred to an experimental tank (30 cm x 30 cm x 10 cm) containing water from the animal’s home tank and respired by a constant flow of oxygenated water through their mouth at a flow rate of 10 mL*min^-1^. Subsequently, the animal’s head was locally anesthetized with lidocaine ointment (5%; AstraZeneca, Mississauga, ON, Canada), the skull was partly exposed, and a small window (∼ 5 mm^2^) was opened over the hindbrain.

### Stimulation

The electric organ discharge of *A. leptorhynchus* is neurogenic, and therefore is not affected by injection of curare. All stimuli consisted of amplitude modulations (AMs or beats) of the animal’s own EOD and were produced by triggering a function generator to emit one cycle of a sine wave for each zero crossing of the EOD as done previously [83]. The frequency of the emitted sine wave was set slightly higher (30 Hz) than that of the EOD, which allowed the output of the function generator to be synchronized to the animal’s discharge. The emitted sine wave was subsequently multiplied with the desired AM waveform (MT3 multiplier; Tucker Davis Technologies), and the resulting signal was isolated from the ground (A395 linear stimulus isolator; World Precision Instruments). The isolated signal was then delivered through a pair of chloridized silver wire electrodes placed 15 cm away from the animal on either side of the recording tank perpendicular to the fish’s rostro-caudal axis (Fig. 1a).

We generated chirp stimulus waveforms (Fig. 1b) with different attributes by systematically varying chirp duration (8, 11, 14, 17, 20 and 29 ms), amplitude (10, 35, 60, 85 and 110 Hz) and phase (0, 45, 90, 135, 180, 225, 270 and 315°) of the underlying beat cycle at which the chirp occurs (Fig. 1c). Importantly, when varying chirp duration or amplitude, we considered chirps occurring at either phase 90° or 270° of the beat cycle with a fixed amplitude of 60 Hz when varying duration, or a fixed duration of 14 ms when varying amplitude, on top of a sinusoidal beat with a frequency of *f*_*beat*_ = 4 Hz, as done previously [25, 32]. Conversely, chirp duration and amplitude were kept constant (14 ms and 60 Hz) when varying phase. As such, we used a total of 30 different chirp waveforms (12 for chirp duration, 10 for chirp amplitude and 8 for chirp phase). Parameter ranges were chosen to contain those observed in previous studies [19, 20, 31, 84]. It is important to note that the chirp amplitude is not equivalent to the actual spectral frequency content of the resulting AM stimulus which is 50-100 Hz [20]. We chose a 4 Hz beat because, as mentioned above, this was the frequency used in previous studies [25, 31]. We note that the chirp stimuli considered here were consistently most frequently produced when the frequency difference between both animals was lowest for various conditions [16, 20, 75, 84]. Previous studies have further demonstrated a sex dimorphism in EOD frequency for *A. leptorhynchus* in that males have significantly higher EOD frequencies than females [85, 86]. As such, same-sex encounters led on average to lower beat frequencies than opposite-sex encounters [22]. To measure the stimulus intensity, a small dipole was placed close to the animal’s skin. Stimulus intensity was adjusted to produce changes in EOD amplitude that were ∼20% of the baseline level, as done previously [25, 31, 34]. Finally, each chirp stimulus waveform (i.e., a chirp with given duration, amplitude or phase) was presented at least 40 times (i.e., 40 trials) with ∼1.5 sec between subsequent presentations, followed by another waveform.

### Recordings

Simultaneous extracellular recordings from both ON- and OFF-type ELL pyramidal cells were made using Neuropixel probes (Imec Inc.). The probe was angled at approximately 15 deg with respect to vertical and advanced between 1500 and 2000 µm into the ELL with reference to the probe tip. Neurons were recorded at depths between 200 and 1400 µm from the brain surface and as such most likely included neurons within the lateral and centrolateral segments [87]. Recordings were digitized at 30 kHz using spikeGLX (Janelia Research Campus) and stored on a hard drive for further analysis. Spikes were sorted using Kilosort [88] and subsequently curated using the phy application (https://github.com/cortex-lab/phy) [30, 33, 89]. Overall, we recorded and sorted from a total of N = 89 ELL pyramidal cells across 4 recording sessions. We note that only neurons with baseline firing rates between 5 and 100 Hz were used for subsequent analyses, leaving a total number of 74 neurons (16 from session 1; 14 from session 2; 27 from session 3 and 17 from session 4).

### Modeling

We modeled electroreceptor afferent filtering properties using the following set of differential equations

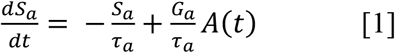

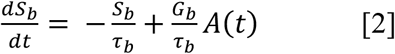

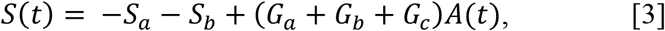

where *A*(*t*) is the stimulus amplitude, *S*(*t*) is the filtered stimulus, *G*_*x*_(*x* = *a, b*, and *c*) are the gain values (spikes/s mV) and *τ*_)_*xx* = *a, b*) are time constants (s). The total synaptic current is given by

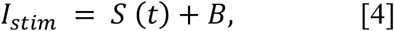

where *I*_*stim*_ is the synaptic current and *B* is a constant.

Next, we developed a two-compartmental biophysical model of ELL pyramidal cell activity [35, 90]. The model consists of two compartments and is based on the Hodgkin-Huxley formalism. The equations describing the membrane voltages in the somatic (*V*_*S*_) and dendritic (*V*_*D*_) compartments are given by

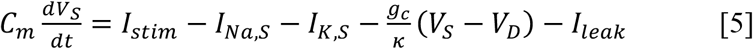

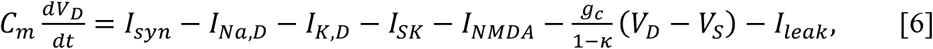

where *C*_*m*_ is the membrane capacitance, *I*_*Na,i*_ and *I*_*K,i*_ are the fast inward sodium and the slow outward delayed rectifier potassium currents in the soma (*i* = *S*) and the dendrite (*i* = *D*), respectively, *I*_*stim*_ is the synaptic current to the somatic compartment from the stimulus and *I*_*syn*_ is synaptic currents to the dendritic compartment, respectively, *I*_*SK*_ and *I*_*NMDA*_ are the outward calcium-dependent small-conductance potassium and the inward NMDA calcium currents acting on the dendrite, and *I*_*leak*_ is the passive leak current present in both compartments. The two compartments are linked together through a resistor, where *g*_*c*_ is the maximum conductance and *κ* is the somatic-to-dendritic area ratio. The currents *I*_*Na,i*_ and *I*_*K,i*_ (*i* = *S, D*) are necessary to generate somatic action potentials and the proper spike backpropagation that yields somatic depolarizing afterpotentials (DAPs) [35]. The two currents *I*_*NMDA*_ and *I*_*SK*_ were added to the dendritic compartment because of the important role they play in regulating spiking activities of ELL pyramidal cells both in vitro [62] and in vivo [61, 91, 92]. The kinetics of the ionic currents included in each compartment (*i* = *S, D*) are given by

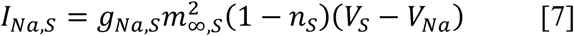

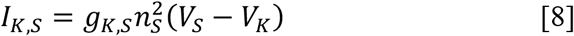

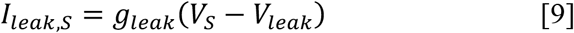

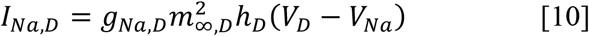

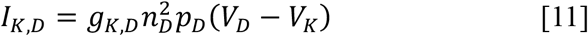

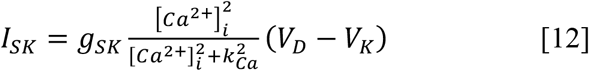

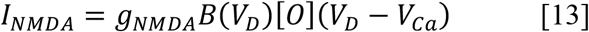

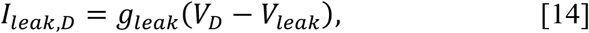

where *g*_*j,i*_ *(j* = *Na, K, leak, SK, NMDA* and *i* = *S, D*) are the maximum conductances, *V*_*j*_ (*j* = *Na, K, Ca, leak*) are the Nernst potentials, [*Ca*^2+^]_*i*_ is the Ca concentration in the dendritic compartment (μM), *k*_*Ca*_ is the half-maximum activation of the SK channel (μM), *B*(*V*_*D*_) is the magnesium block, given by [93]

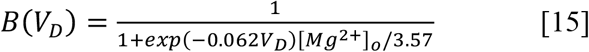

[*Mg*^*2+*^]_*o*_ is the extracellular magnesium concentration (mM), [*O*] is the open probability of the NMDA receptors defined by a Markov model comprised of three closed, one open and one desensitized states with transition rates *R*_*i*_ (*i* = *b, u, c, o, d, r*) between them, *m*_∞,*i*_ *(i* = *S, D)* is the steady state activation of *I*_*Na,i*_, and *x (x* = *n*_*i*_, *h*_*D*_, *p*_*D*_; *i* = *S, D)* are the activation/inactivation gating variables whose steady states and time constants are denoted by *x*_∞_ and *τ*_*x*_, respectively, and whose dynamics are governed by the equation

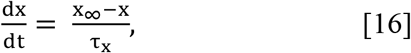

The steady state activation/inactivation functions are given by

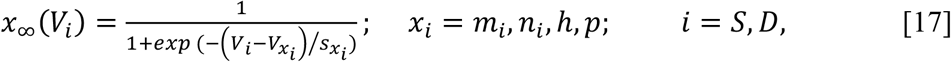

In this model formalism, it was assumed that *I*_*SK*_ and *I*_*NMDA*_ are co-localized within spines. We therefore did not consider calcium diffusion within/between spines.

The NMDA receptor activation depends on the concentration of glutamate following principles of ligand-gated channels described previously [93], where the transition rates between unbound and bound states is based on the concentration of ligand. In our model, the timing of the release of glutamate follows a Poisson process formalism with firing rate *λ*_*glu*_. The glutamate concentration following each release event was described by an alpha function [93], given by

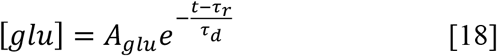

where *τ*_*r*_ is the timing of glutamate release, determined by presynaptic spike times, and *τ*_*d*_ is the time constant of the alpha function, and *A*_*glu*_ is a scaling constant. The parameter *τ*_*d*_ was chosen to be fast enough to prevent glutamate release events from overlapping.

The calcium mobilization model followed the flux-balance formalism; it describes fluxes across the cell and ER membranes in the dendritic compartment. Calcium mobilization across the cell membrane includes three fluxes through the NMDA receptors: *J*_*NMDA*_ = *αI*_*NMDA*_, where 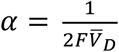(*F* is Faraday’s constant and 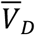 is the volume of the dendritic compartment), plasma membrane calcium ATPases (PMCA) pumps: *J*_*PMCA*_, and leak: *J*_*INLeak*_. Calcium mobilization across the ER membrane, on the other hand, includes three fluxes through the IP3Rs: J_IP3R_, sarco/endoplasmic reticulum ATPases (SERCA) pumps: J_SERCA_, and leak: *J*_*ERLeak*_. The Li-Rinzel model was adopted to describe IP3R kinetics [94, 95]. It is given by

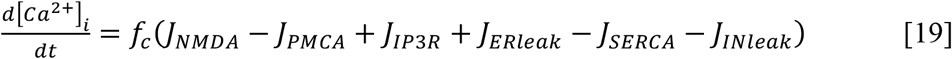

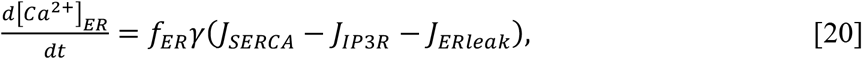

where *f*_*c*_ (*f*_*ER*_) is the fraction of free calcium concentration in the cytoslic (ER) component of the dendrite, and γ is the volume ratio of cytosol to ER in the dendrite. The fluxes through the PMCA and SERCA pumps are given by

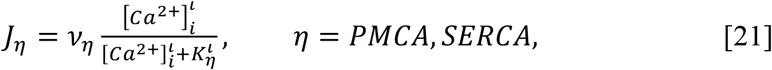

where *ι* = 2 is the Hill coefficient, *ν*_*η*_ are the maximum flux rates (μM/s), and *K*_*η*_ are the half-maximum activations for calcium flux (μM). Fluxes due to leak across cell and ER membranes are given by

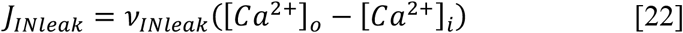

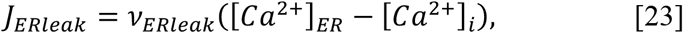

where *ν*_*ξ*_ (*ξ* = *INleak, ERleak*) are the maximum flux rates (μM/s). Finally, flux through IP3Rs is adopted from the Li-Rinzel model [94] and is given by

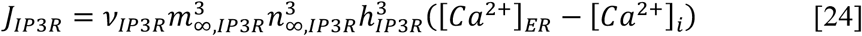

where *ν*_*IP*3*R*_ is the maximum flux rate (μM/s),

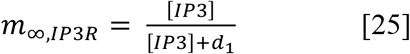

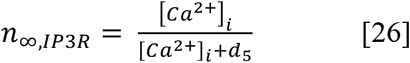

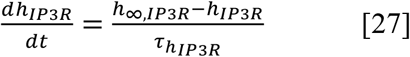

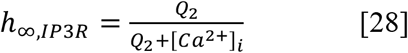

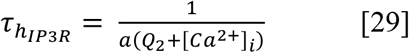

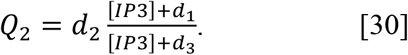

Note that [*IP*3] is the cytosolic concentration of IP3 in the dendrite (μM).

The synaptic input current, *I*_*syn*_, was used to represent all the stochastic background excitatory and inhibitory pre-synaptic activity, defined by *W*_*x*_(*t*) (*x* = *E, I*), applied to the dendritic compartment [96, 97]. It has been previously suggested that the power spectra of excitatory and inhibitory background activity in synaptic inputs exhibit a power law spectrum, *S*_*x*_(*f*)∼ 1/*f*^*β*^, where *β* determines the exponent of steepness of the slope of the power spectrum [97]. Based on this, the total synaptic input incorporated into the model can be described by

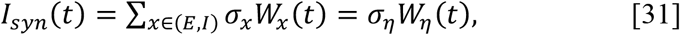

where *σ*_*η*_ is the total noise intensity obtained by integrating the two components, *W*_*η*_ (*t*) is the sum of both excitatory and inhibitory pre-synaptic inputs reconstructed using the following expression

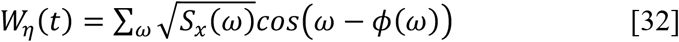

with *ω* = 2*πf* and *ϕ*(*ω*) ∈ [0,2*π*] is a random phase multiplied by the spectrum in the frequency domain. Once the spectrum is reconstructed, the synaptic time series can be obtained using inverse Fourier transform of the spectrum as described previously [98]. Model simulations were performed in python using the Euler-Maruyama method [99] as the integration solver of the full stochastic model. The excitatory and inhibitory pre-synaptic inputs were generated using the algorithm described previously [98]. Initially, we ran all simulations using an integration timestep of 0.01 ms (with a 100 kHz sampling rate). The membrane voltage trace was then downsampled to 20 kHz.

### Data Analysis

All data analysis was performed offline using custom written codes in Matlab software (MathWorks). Sorted and curated spike times for each neuron were converted into “binary” sequences sampled at 2 kHz. The content of a given bin was set to 2000 if a spike occurred within it and 0 otherwise. We note that, as the binwidth was smaller than the refractory period of ELL pyramidal cells, there can be at most one spike occurring during any given bin. The time-dependent firing rate of neuron *i, X*_*i*_(*t*), was obtained by filtering the binary sequence using a box-car filter with 6 ms. Results were normalized by the maximum value, such that the PSTH ranges between 0 and 1.

### Separating burst and isolated spikes

We used an interspike interval (ISI) threshold to assign each spike as part of a burst or not (i.e. “isolated”), and thus generated the burst spike trains and the complementary isolated spike trains [43, 44, 64, 68, 70, 100] (sub. Fig. 1a). Specifically, spikes with interspike interval lower or equal than the threshold were assigned as part of a burst with the process continuing until an ISI is greater than the threshold. Spikes that were not part of bursts were deemed to be isolated. The burst fraction was defined as the fraction of ISIs that are less than or equal to a given threshold value. Threshold values were varied between 0.1 and 40 ms with 30 bins spaced logarithmically (sub. Figs. 1b, c). The main figures show results obtained using burst threshold values chosen from the trough of the bimodal ISI distribution.

### Computing response correlations

We quantified response correlations using the Pearson correlation coefficient (function “corrcoef” in Matlab) between the filtered activities from neurons of our largest dataset (n = 27) as well as from our model neurons (n = 27) to all chirp stimuli during a 100 ms evaluation time window. The waveforms were shifted to the right by 7 ms to account for axonal and synaptic transmission delays [81]. This value was chosen because it comprises the neural response for most of the stimuli considered. Furthermore, we also quantified response correlations between the filtered activities from all neurons (n = 74) pooled across all recording sessions to test the effect of a larger population, as we saw that noise correlations did not affect our results significantly (sup. Fig. 2). In all cases, we considered the full spike trains, burst spike trains, or isolated spike trains, respectively. We computed the correlations between all stimulus waveforms themselves using 100 ms of the stimulus waveform following chirp onset.

### Computing noise correlations

We quantified noise correlations between neuron activities from all neurons (n = 74) recorded during all four recording sessions using the time dependent firing rates that were obtained from each binary spike train. We then computed the correlation coefficient between the two firing rate sequences using Pearson’s correlation coefficient

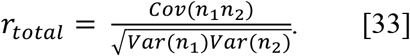

Noise correlations were computed as the correlation coefficient between the firing rate residual sequences. The firing rate sequences were first averaged over trials and the mean firing rate sequence (i.e., that due to the stimulus) was then subtracted from the firing rate for each trial to obtain the residual firing rate sequences. As such, the residual firing rate sequences represent the component of the neural response that cannot be explained by a given stimulus waveform (i.e., “noise”) because, unlike the stimulus waveform itself, these are not constant across trials.

### Classifier

We used a classifier to quantify the performance of ELL pyramidal cell populations at stimulus discrimination when considering the full spike trains, burst spike trains or isolated spike trains. For each individual chirp stimulus, the combined response to a random trial was taken as the template for a given chirp stimulus waveform. Next, each combined response was assigned as being generated by the stimulus that gave rise to a given template based on whether the distance between the combined response and the template was minimum. We thus constructed a “confusion matrix” whose element (*i,j*) gives the probability that a response was assigned as being generated by stimulus *j* given that it was actually generated by stimulus *i* [33, 47, 48]. The diagonal elements of this matrix are the probabilities that a stimulus was correctly assigned, whereas non-zero off-diagonal elements indicate misclassification. For each confusion matrix obtained from the metric-space analysis, we computed the discrimination performance by averaging over the diagonal elements. The discrimination performance thus varied between 0 (no discrimination) and 1 (perfect discrimination). Note that the chance level for discrimination performance was ∼0.03 (that is, 1/30) because we used a total of 30 different chirp stimuli. The distance between combined neuron activities was computed using the van Rossum metric [101]. First, the combined neuronal activities were convolved with a decaying exponential kernel with time constant *τ*

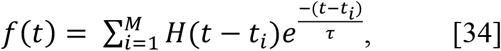

where *t*_*i*_ is the *i*^*th*^ spike time, *M* is the total number of spikes and *H(t)* is the Heaviside step function (*H(x)* = 0 if x<0 and *H(x)* = 1 if x >= 0) and *τ* =3 ms [30]. Next, the van Rossum distance was computed as the Euclidian distance between convolved combined neuronal activities *fR*_*j*_ and *fR*_*K*_ [102]

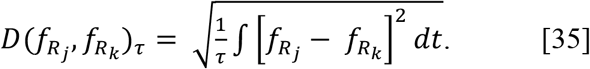

Performances reported in the paper were averaged over templates and error bars were generated over the total number of simulations.

### Statistics

Values were reported as mean ∓ STD. Statistical tests were performed using a one-way ANOVA with Bonferroni correction unless otherwise stated.

## Figure captions

**Supplementary Figure 1:**
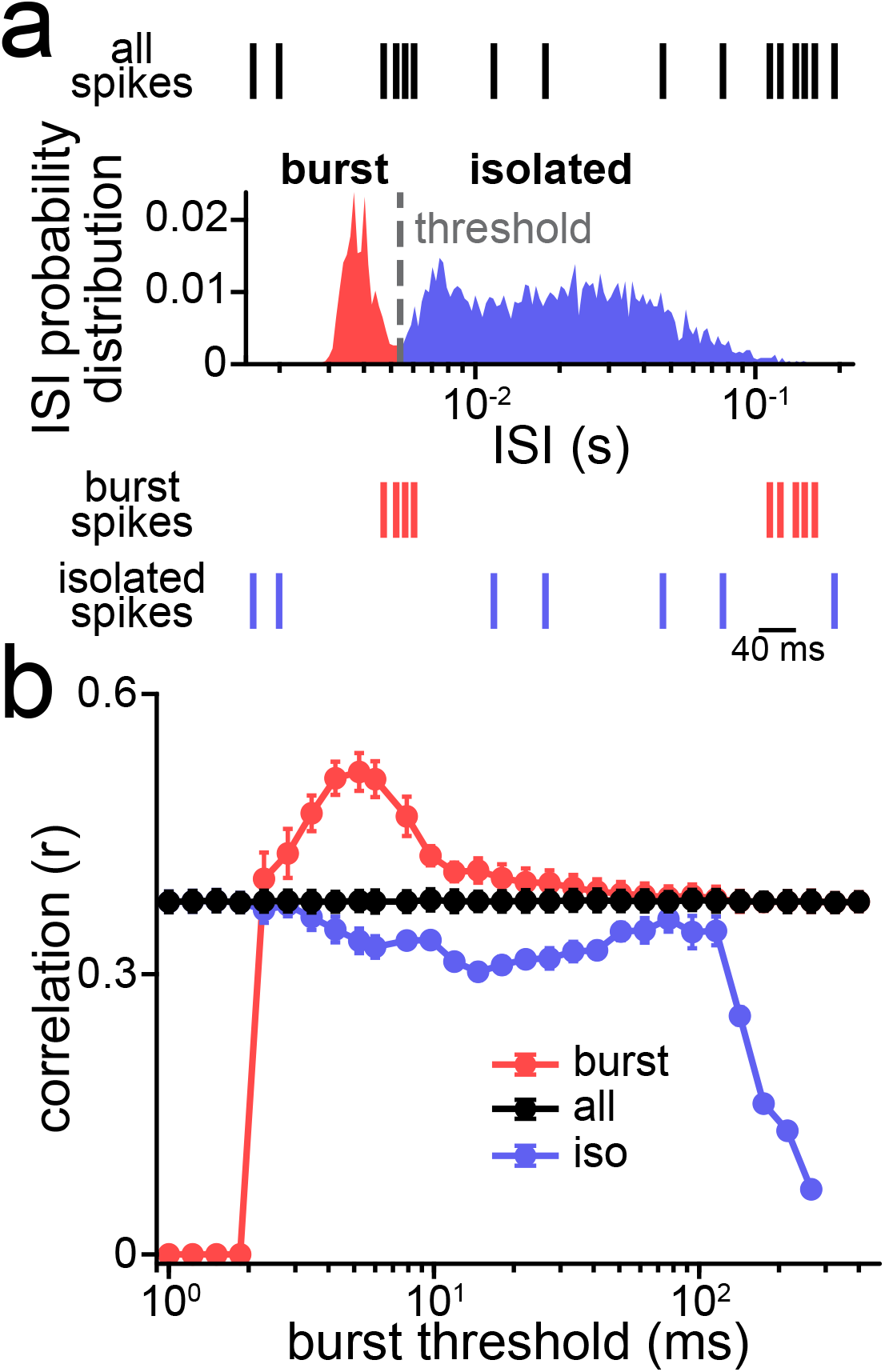
ELL pyramidal cells display burst firing. **(a)** ISI distribution from an example ELL pyramidal cell was bimodal (Hartigan’s dip test; p = 1*10^−3^). The trough of the distribution (dashed line) was used to separate the full spike train (i.e., all spikes, top, black) into spikes that belong to bursts (i.e., the “burst spike train”, red) and those that did not (i.e., the “isolated spike train”, blue). **(b)** Correlation as a function of burst threshold obtained for the full (black), burst (red) and isolated (blue) spike trains.

**Supplementary Figure 2:**
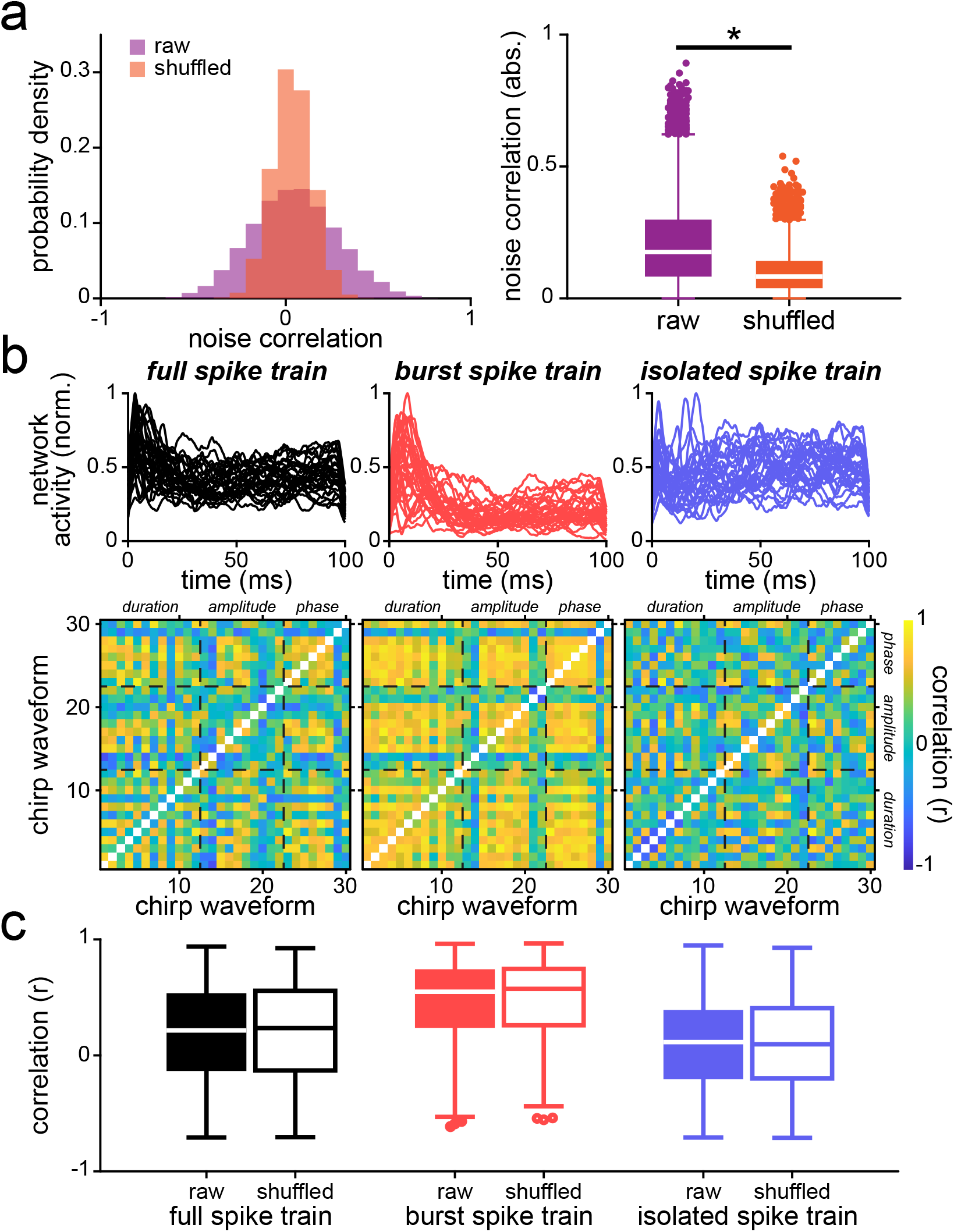
Noise correlations do not influence population coding of natural electrocommunication stimuli. **(a)** *Left*: Probability density functions of noise correlations in a raw dataset (n = 27; orange) and after shuffling the trials (purple). *Right*: Noise correlation magnitude significantly decreases after randomly shuffling responses during repeated presentations of the stimulus (paired t-test; p < 0.001). **(b)** *Top*: Superimposed network activity in response to all chirp stimulus waveforms obtained for the full (black, left), burst (red, middle) and isolated spike trains (blue, right) when noise correlations were removed. *Bottom*: Correlation matrices of network activities when considering the full (left), burst (middle) and isolated spike trains (right). **(c)** Whisker-box plots of correlation coefficients obtained for network activities without (solid) and with (hollow) noise correlations when considering either the full (black), burst (red) and isolated (blue) spike trains. No significant differences were observed (one-way ANOVA; df = 5; F = 78.36; p >> 0.1; Bonferroni corrected).

**Supplementary Figure 3:**
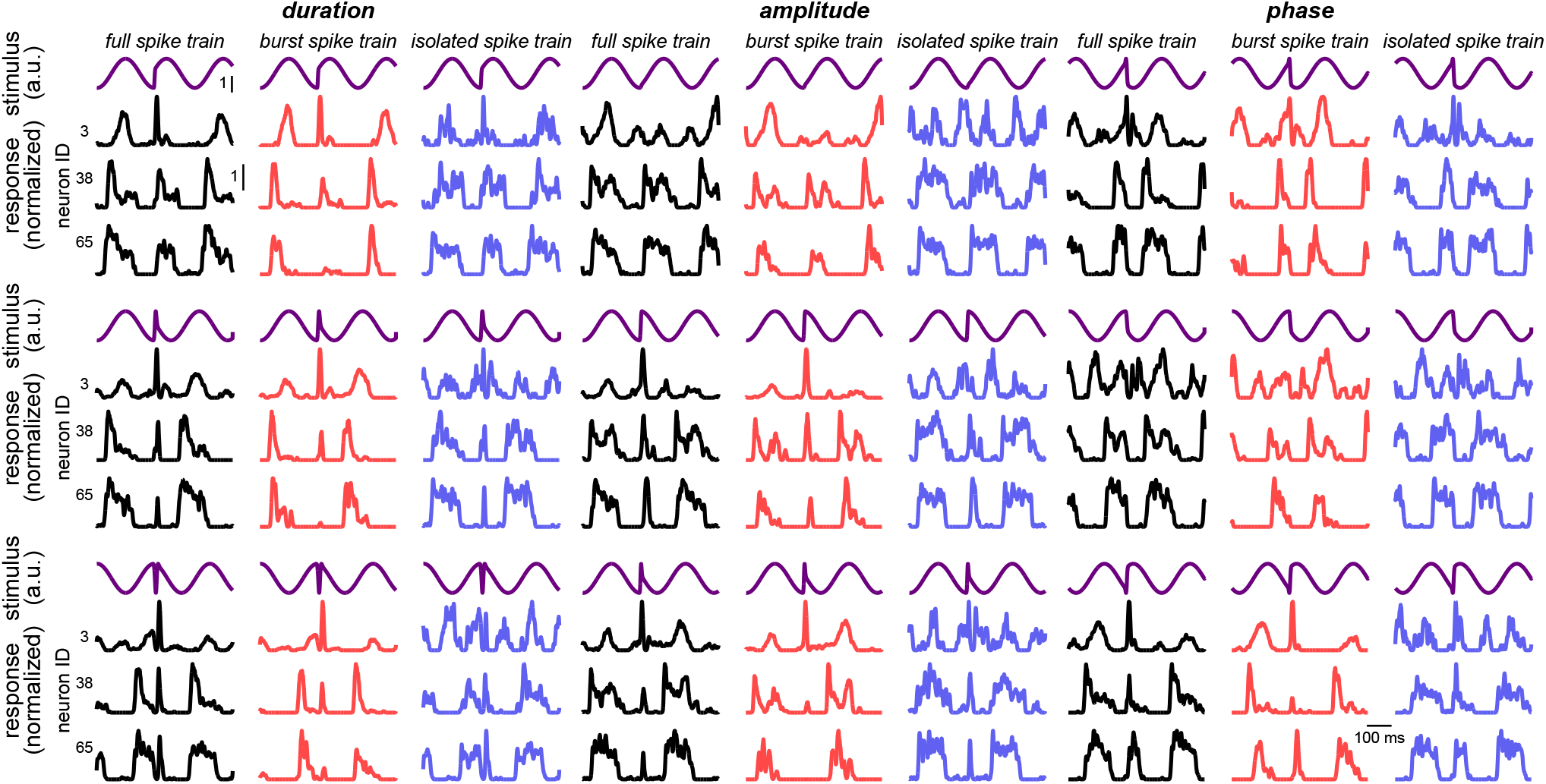
Examples network activities in response to different chirp waveforms. **(a)** Network activity in response to stimulus waveforms when varying chirp duration (top left, purple), chirp amplitude (top middle, purple) and chirp phase (top right, purple) when considering the full (i.e., all spikes, black; middle top), burst (red, middle bottom) and isolated (blue, bottom) spike trains. Note that the responses of the burst spike trains are overall more similar during the chirp window for different chirp waveforms than either all or isolated spike trains.

**Supplementary Figure 4:**
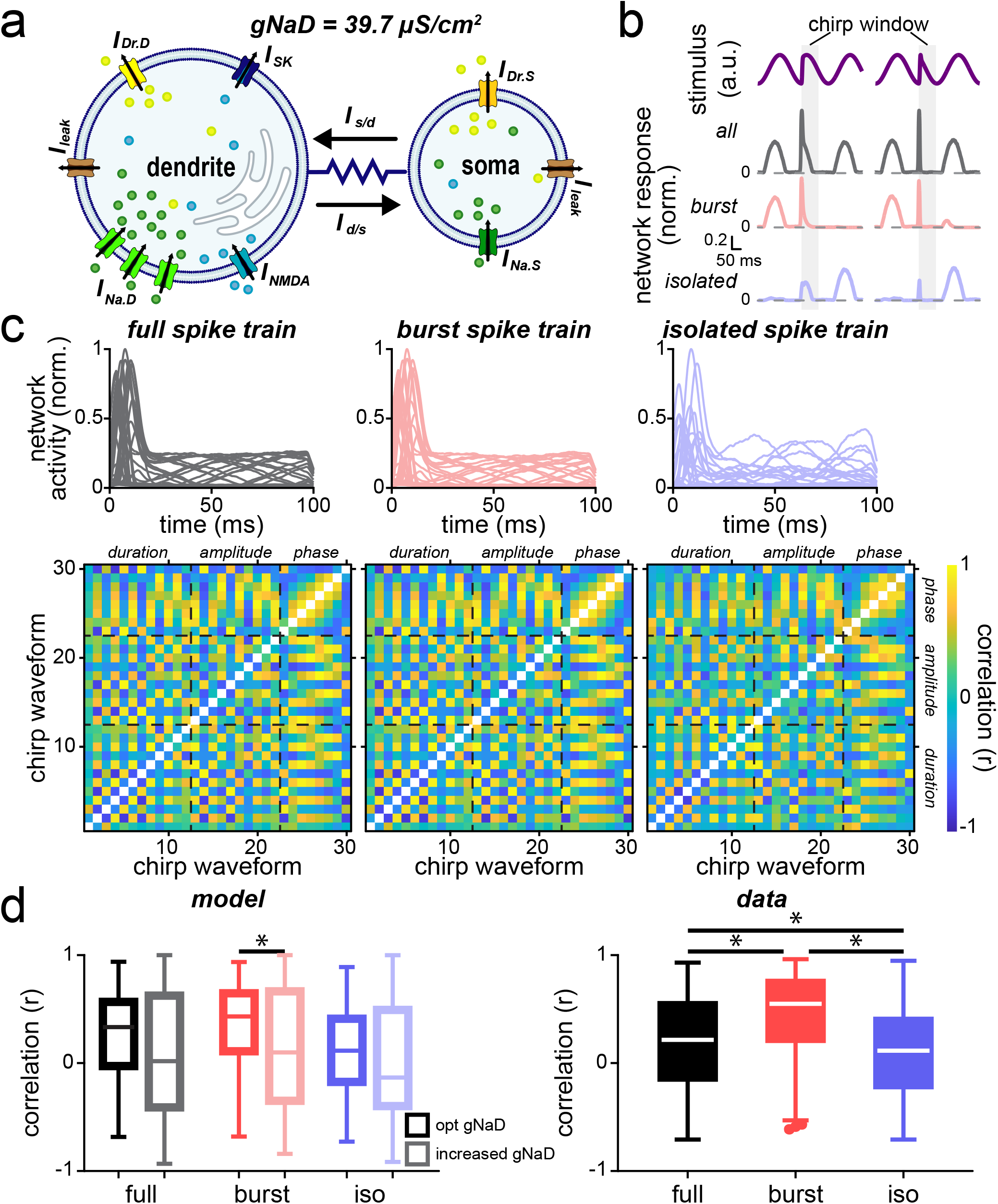
An active burst mechanism enhances invariant population coding of natural electrocommunication stimuli. **(a)** Schematic of the model with a low *g*_*NaD*_ value of 1.6 μS/cm^2^. **(b)** Examples of model network activities in response to two chirp waveforms (top, purple) using the full (i.e., all spikes, black; middle top), burst (red, middle bottom) and isolated (blue, bottom) spike trains, respectively. **(c)** *Top*: Superimposed network activity traces from the model in response to all chirp stimuli when considering either the full (black, left), burst (red, middle) and isolated spike trains (blue, right). *Bottom*: Correlation matrices of network activities when considering the full (left), burst (middle) and isolated spike trains (right). **(d)** Whisker-box plots of correlation coefficients obtained for network activities associated with the: model with optimal value of *g*_*NaD*_ (left, hollow) and model with low *g*_*NaD*_ value (left, lighter shades) and experimental data (right, solid) when considering either the full (black), burst (red) and isolated (blue) spike trains. Correlation for bursts in the model was significantly less for low values of *g*_*NaD*_ (one-way ANOVA, df = 5; F = 24.00; p_full_ = 0.99; p_burst_ = 1.98*10^−14^; p_iso_ = 5.85*10^−4^; Bonferroni corrected).

**Supplementary Figure 5:**
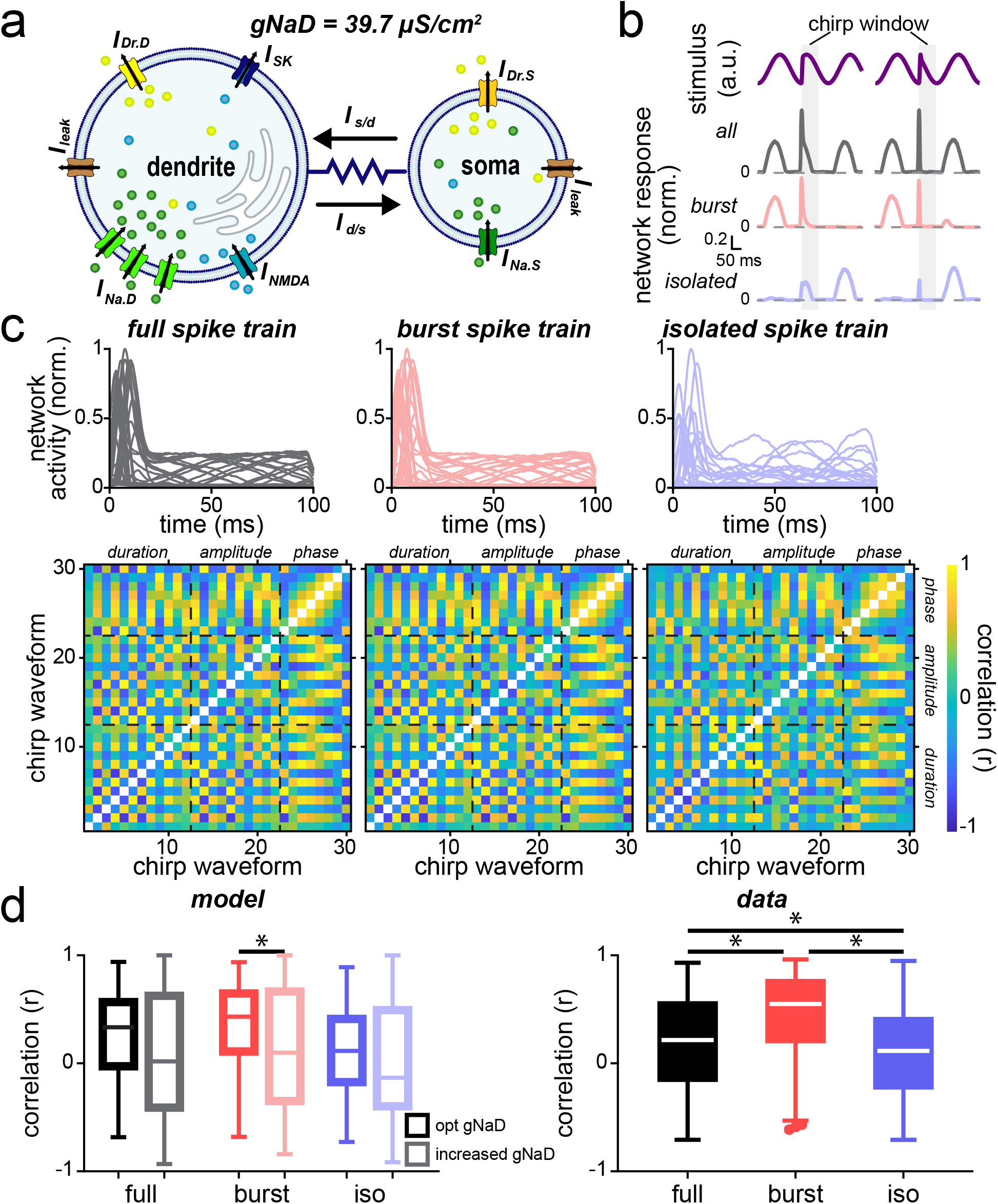
Large levels of burst firing are detrimental to invariant population coding of natural electrocommunication stimuli. **(a)** Schematic of the model with a high *g*_*NaD*_ value of 39.7 μS/cm^2^. **(b)** Examples of model network activities in response to two chirp waveforms (top, purple) using the full (i.e., all spikes, black; middle top), burst (red, middle bottom) and isolated (blue, bottom) spike trains, respectively. **(c)** *Top*: Superimposed network activity traces from the model in response to all chirp stimuli when considering either the full (black, left), burst (red, middle) and isolated spike trains (blue, right). *Bottom*: Correlation matrices of network activities when considering the full (left), burst (middle) and isolated spike trains (right). **(d)** Whisker-box plots of correlation coefficients obtained for the network activities associated with the: model with optimal value of *g*_*NaD*_ (left, hollow) and model with high *g*_*NaD*_ value (left, lighter shades) and experimental data (right, solid) when considering either the full (black), burst (red) and isolated (blue) spike trains. Correlation for bursts in the model was significantly less for high values of *g*_*NaD*_ (one-way ANOVA, df = 5; F = 27.00; p_full_ = 1.20*10^−6^; p_burst_ = 1.63*10^−10^; p_iso_ = 0.44; Bonferroni corrected).

## Notes

### Competing Interest Statement

The authors have declared no competing interest.

